# Determination and characterization of papillomaviurs and parvovirus causing mass mortality of Chinese tongue sole (*Cynoglossus semilaevis*) in China

**DOI:** 10.1101/2024.01.02.573988

**Authors:** Shuxia Xue, Xinrui Liu, Yuru Liu, Chang Lu, Lei Jia, Yanguang Yu, Houfu Liu, Siyu Yang, Zhu Zeng, Hui Li, Jiatong Qin, Yuxuan Wang, Jinsheng Sun

**Author notes:** Correspondence: Jinsheng Sun, The key Laboratory of Animal and Plant Resistance / College of Life Science, Tianjin Normal University, Tianjin, 300387, China.

## Abstract

Chinese tongue sole (*Cynoglossus semilaevis*) is one of the representative species in flatfish aquaculture in China. In recent years, massive mortality of farmed Chinese tongue soles occurred in Tianjin, China. The causative pathogens were determined as *Cynoglossus semilaevis* papillomavirus (CsPaV) and parvovirus (CsPV) by electron microscopy, virus isolation, experimental challenge, histopathological analysis, genome sequencing, fluorescence *In situ* hybridization (FISH) and epidemiology investigation. Electron microscopy showed large numbers of spherical non-enveloped virus particles presenting in liver, kidney, spleen, gill and heart of the diseased fish. The viruses were isolated and propagated in flounder gill cells (FG) and induced a typical cytopathic effect (CPE). The cumulative mortality reached 100% at 8 dpi by intraperitoneal injection. The complete genome of CsPaV and CsPV was 5939 bp and 3663 bp in size respectively, and both viral genomes shared no nucleotide sequence similarity with other viruses. The CsPaV contained seven predicated protein coding regions (E1, E2, L2, L1a, L1b, sORF1 and sORF2) and CsPV contained three predicated protein coding regions (NS1, VP and ORF3). Phylogenetic analysis basing on L1 and NS1 protein sequences revealed that CsPaV and CsPV are novel members belonging to new genus in *Papillomaviridae* and *Parvoviridae* family. FISH results showed positive signals in spleen and kidney tissues from CsPaV and CsPV infected fish and the two viruses could co-infected one cell. Epidemiological investigation showed that the two viruses cocurrented in 82.9% sampled fish and they were proved to be the pathogenic agents of the emerging disease in farmed Chinese tongue soles in China. This study represents the first report of co-infection of papillomavirus and parvovirus in farmed fish and provides a basis for further studies on prevention and treatment of the emerging viral disease, and also represents clues to elucidate the the mechanisms of viruses co-infection and evolution of viruses.

**Author summary:** Chinese tongue sole is a valuable fish kept in marincultures. Outbreak of an emerging disease caused massive mortality and resulted in significant economic loss. The pathogenic agent remains unknown. In this study, we identified papillomavirus (CsPaV) and parvovirus (CsPV) from the diseased fish simultaneously, and they are proved to be novel members belonging to new genus in *Papillomaviridae* and *Parvoviridae* family. It was shown that the emerging disease was caused by co-infection with the two viruses. Viral co-infections are widespread in nature, however, studies and available data on viral co-infections in fish aquaculture are limited. Our findings represent new clues to elucidate the mechanisms of viruses co-infection and evolution of viruses, and moreover, the present study provide a solution for the control of emerging viral diseases in Chinese tongue sole.

## Introduction

Chinese tongue soles (*Cynoglossus semilaevis*) are naturally distributed in the offshore waters of the Yellow Sea and Bohai Sea, belonging to the family *Soleidae* of the order *Flounder*. With the breakthrough development of artificial broodstock technology, the breeding scale of Chinese tongue sole has gradually expanded, and it becomes an important marincultured species in China. The production of the fish equals 2,500-3,000 tons in 2021 with a current value excess 500 million dollars [1].

Previous studies showed that *vibrios* were the most common pathogens in the production of Chinese tongue soles, and researches on vibirosis have prevented the epidemic of the disease [2–4]. In recent years, emerging viral diseases have threatened the aquaculture of Chinese tongue soles. However, reports of viral diseases associated with Chinese tongue soles are limited. A betanodavirus have been isolated from diseased Chinese tongue sole in China and its genome sequence shares ≥ 98.7% similarity with grouper nervous necrosis [5]. Xiao et al. (2015) [6] have reported that a viral pathogen was responsible for the development of splenic necrosis sings in tongue sole, but no further studies have been elucidated. In December 2021, massive mortality of Chinese tongue soles occurred in a farm in Tianjin, China. No parasites were examined from the diseased fish. *Vibrios* were found in partially diseased fish, but they are not associated with the clinical signs observed in the study. A large number of virus particles shaped in spherical were observed in the tissues of the diseased fish by electron microscopy. No reported viral pathogens were detected based on routine diagnostic methods. These results suggested that novel viral pathogen was probably responsible for the emerging disease of Chinese tongue soles.

In the present study, co-infection of two viruses determined as papillomavirus (named CsPaV) and parvovirus (named CsPV) was proved to be responsible for the emerging disease of Chinese tongue soles. The results of the study provide a basis for further study on the co-infection mechanism of the two viruses and the prevention and control method for the disease in future.

## Materials and methods

### Fish samples

Diseased Chinese tongue soles with body length 37-40 cm were collected from a commercial farm in Tianjin, China in December 2021. The diseased fish were transferred alive in oxygenated bags to laboratory for diagnosis and detection of the pathogens. Healthy Chinese tongue soles with body length 25-30cm for pathogen challenge assay were obtained from another farm in Tianjin and no history of disease was recorded. All fish were maintained in fiberglass tanks at 21 °C, 21‰ salinity with aeration for 1 week and no diseased syndromes were shown before experimental challenge.

### Electron microscopy

Liver, kidney, spleen, gill, heart and brain tissues of naturally diseased fish were fixed in 2.5% glutaraldehyde and a 1% osmium tetroxide for post-fixation, then dehydrated, embed, cut and stained with 2% uranyl acetate and lead citrate. All samples were then observed with a transmission electron microscope at 80 KV (JEM1200, Janpan).

### Cell culture and virus isolation

FG cells were cultured in DMEM medium (GIBCO, USA) supplemented with 10% fetal bovine serum (FBS, GIBCO, USA) at 23 °C in 2.5% CO_2_ atmosphere. The mixed tissues of liver, kidney and spleen from naturally diseased Chinese tongue soles were grinded with liquid nitrogen and homogenized with 10 folds volume of DMEM medium. The suspension was freeze-thawed for 3 cycles (−80 °C to room temperature), centrifuged at 12,300 g for 20 min at 4 °C and then filtered through 0.22 μm filter membrane, and finally, 1 mL of filtrate at 1:10. 1:100 and 1:1000 dilutions were inoculated onto confluent monolayer of FG cells in a 25 cm^2^ flask (Corning, USA). The negative (mock infection) control was inoculated with the same volume of DMED medium. After absorption for 1 hours at 23 °C, 5 mL DMEM medium (containing 2% FBS, 100 IU/mL penicillin, 100 μg/mL streptomycin) was added to the flasks. The cells cultures were incubated at 23 °C and checked daily under an inverted phase contrast microscope (Nikon, Japan) to observe the cytopathic effect (CPE). The supernatant of cells showing obvious CPE was harvested following a 3 times freeze-thaw cycle and centrifugated at 3000 × g for 20 min. Subculture of the isolated virus using methods described above. The virus titer was determined using the 50% tissue culture infective dose (TCID_50_) method in a 96-well culture plate.

To determine the replication dynamics of the viruses in FG cells, the subculture at the first passage at 8 dpi was harvested and used to infect FG cells. TaqMan probe-based quantitative PCR was used to analyze virus copy numbers at 12, 24, 48, 60, 72, 84, 96, 108, 120, 132, 144, 156, 168 and 180 hours post-infection. The viral genomic DNA was extracted from the infected cells and supernatant using DNA viral kit (Omega, Germany) according to the manufacturer’s instructions. Quantitative PCR was conducted as described mentioned below.

### Histopathological observation

Tissues samples of liver, spleen, kidney and intestine from experimentally infected fish for 3, 5 and 7 days were collected for histopathological observation by using hematoxylin and eosin (HE) staining. Tissues were fixed in 4% paraformaldehyde for 24 hours at 4 °C and washed with Dulbecco’s phosphate-buffered saline (DPBS, Sigma, USA). After dehydrated through a graded ethanol series to absolute ethanol, optimum cutting temperature compound (OCT) was used to embed the samples, which were then cut into 8-μm-thick sections using a cryostat (RM2016, Leica, Germany). The slices were stained with HE and examined by light microscopy (Nikon Eclipse E100, Japan).

### Viral genome sequencing and PCR assays

The mixed tissues of liver, kidney and spleen of diseased fish were collected, homogenized and centrifuged to remove cellular debris, and the supernatant processed by a 0.22 μm filter. The virus was enriched by ultracentrifugation (Beckman Coulter Optima™ L-80XP) at 246,347g for 4 hours at 4 °C. The fish nucleic acids were degraded by 200 units Benzonase (Sigma, USA), 6 units of Turbo™ DNase (Invitrogen, USA) and 15 units RNase Ⅰ (Invitrogen, USA) incubating at 37 °C for 1h. RNA and DNA were extracted using MiniBEST viral RNA/DNA extraction kit (Takara, Japan) according to manufacturer’s instructions, respectively. Sequence independent single primer amplification (SISPA-PCR) method was used to analyze the sequences of the unknown virus [7]. Briefly, viral RNA was converted to cDNA using FR26RV-N (Table 1), and then the viral DNA and the cDNA were processed with Klenow polymerase (Takara, Japan) in the presence of FR26RV-N, finally the double stranded DNA was amplified by PCR using the primer FR20RV (Table 1). The PCR products were used to constructed libraries using NEB Next Ultra DNA Library Prep Kit for Illumina (NEB, USA) following manufacturer’s recommendations. Subsequently, the qualified libraries were pooled and sequenced on Illumina platforms with PE150 strategy according to effective library concentration and data amount required. The residual contaminated nucleic acids were filtered by blast with genome of *Cynoglossus semilaevis* [8] using Bowtie 2 software. The filtered reads were assembled into large scaffolds using SPAdes software. Similartiy searches were performed using Blast X to find the similar matches with other virus in the GeneBank database. The final genomic sequences were obtained by mapping filtered reads against the scaffolds using Soap *denovo* 2.0. Multiple primers (Table 1) were designed to amplify the virus genomes and the genomic sequences were confirmed by Sanger sequencing. The predicted ORFs were analyzed using ORF finder, and the circular genome map of CsPpV was drawn with DNA Plotter. The splicing sites were predicted using Net-Gene2 Server (http://www.cbs.dtu.dk/services/NetGene2/).

**Table 1.**
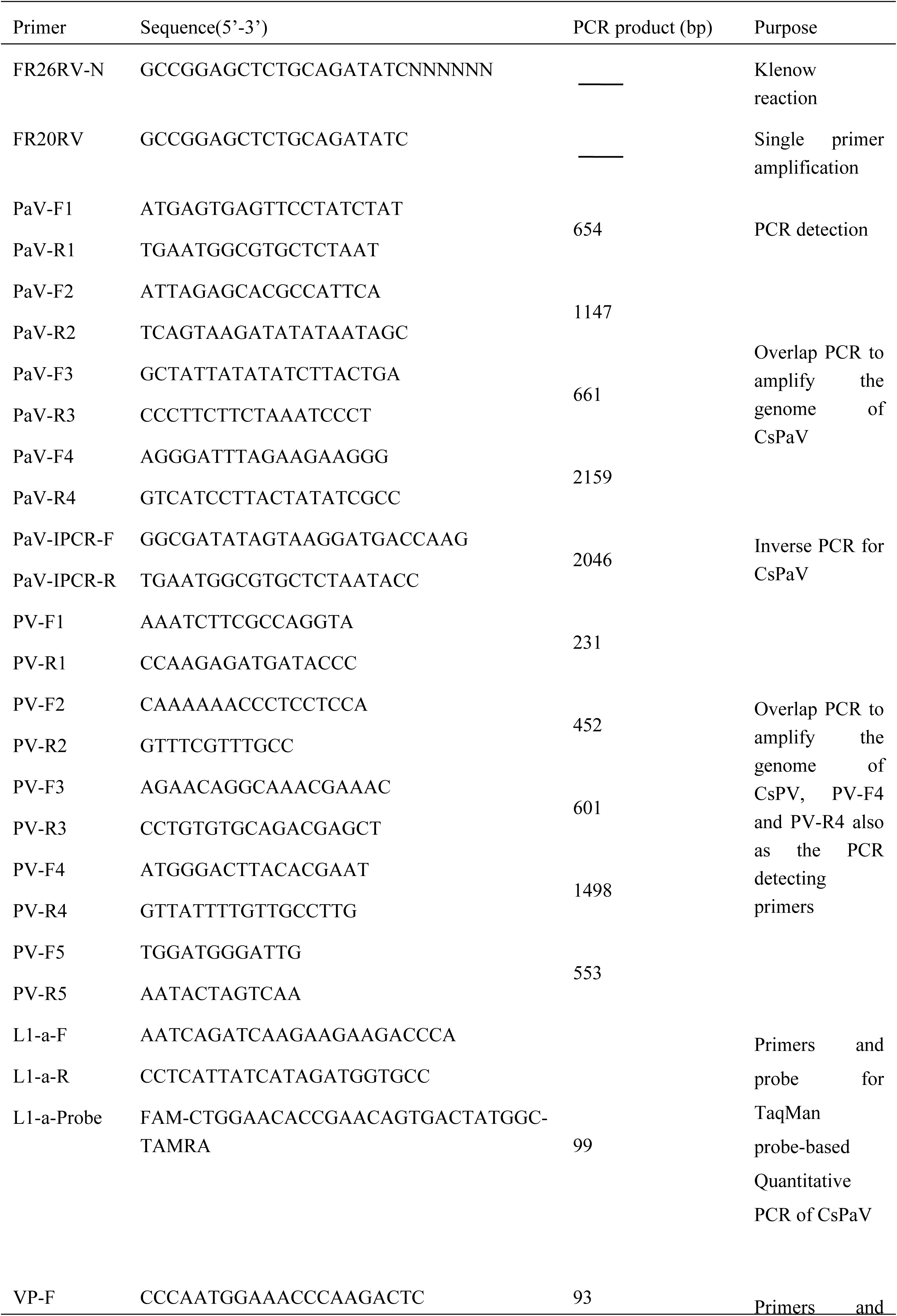
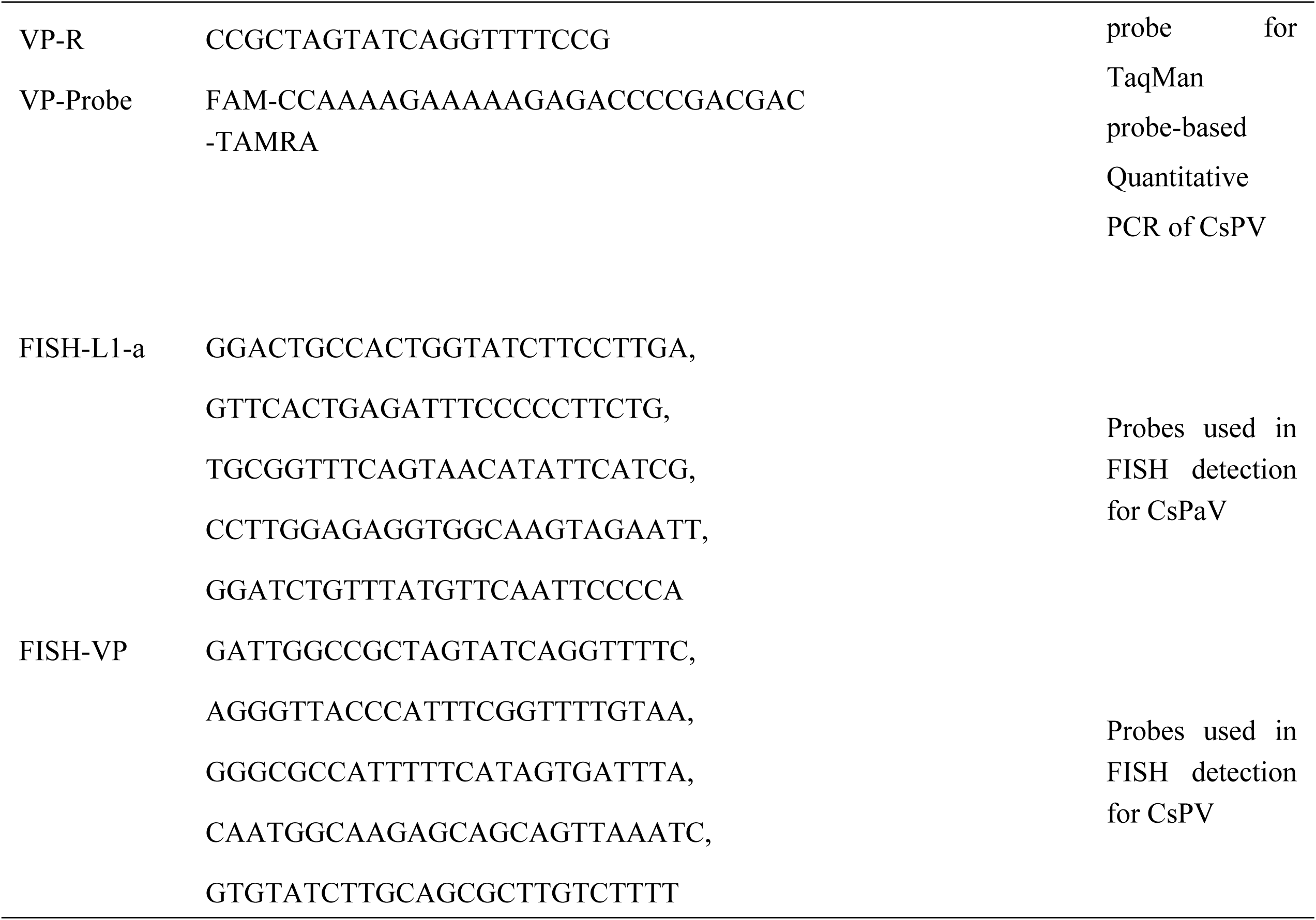
Primers and probes used in the present study.

### RNA extraction and L1 gene sequencing

The complete L1 gene was obtained from genome DNA of CsPaV and inserted into pEGFP-C1, constructing plasmid pEGFP-C1-L1. Transfections were performed using lipo3000 according to manufacturer’s instructions. Total RNA was extracted from viral infected tissues, FG cells and transfected FG cells using TRIzol Reagent. First-strand cDNA synthesis was performed using M-MLV Reverse Transcriptase (Promega) and specific primer of L1 gene. And then the cDNA samples were subjected to PCR and PCR products were sequenced and blast with L1 gene.

### Phylogenetic analysis

Similarity comparison was conducted using BLASTn and BLASTp (NCBI database). The deduced amino acid sequences were analyzed using the Expasy (https://web.expasy.org/translate/). Multiple sequence alignments were performed using Muscle package with default parameters. The phylogenetic trees were constructed basing on L1 nucleotide sequence with other members of family *Papillomaviridae* (http://pave.niaid.nih.gov), L1 amino acid sequences of the identified fish papillomavirus, and NS1 amino acid sequences from other representatives of subfamily *Parvovirinae* and *HamaParvovirinae* with MEGA 10.0 by Maximum-likehood method with the substitution model of Poisson and rates of Gamma Distributed. The node was obtained with 100 bootstrap iterations.

### Fluorescence *In situ* hybridization (FISH)

To investigate the location of mRNAs of CsPaV and CsPV in the infected Chinese tongue soles, Paraffin-SweAMI-double FISH method was performed with FAM-conjugated probes for the CsPaV L1 and Cy3-conjugated probes for CsPV VP gene sequence (Table 1). Liver and spleen samples from experimentally infected fish for 3 days were fixed in 4% paraformaldehyde (PFA, Sigma, USA), embedded in paraffin and sectioned (4 μm) at −20°C (RM2016, Leica, Germany).

The sections were roasted at 62 °C for 2 hours followed by dewaxing and dehydration with ethanol, then digested with proteinase K (20 μg/ml). The sections were hybridized overnight at 40 °C with conjugated probes. After washing with the hybridization solution, the corresponding branch probes were added for hybridization. Then, the signal probes at the dilution ratio of 1: 400 were added and hybridized overnight at 40 °C. Next, DAPI solution was dripped into the sections to counter stain the nucleus. DAPI glows blue by UV excitation wavelength 330-380 nm and emission wavelength 420 nm; FAM glows green (L1 probe of CsPaV) by excitation wavelength 465-495 nm and emission wavelength 515-555 nm; CY3 glows red (VP probe of CsPV) by excitation wavelength 510-560 nm and emission wavelength 590 nm. All the sections were examined by inverted fluorescence microscope (Nikon Eclipse TI-SR, Japan).

### PCR detection

DNA was extracted by DNA viral kit (Omega, Germany) according to the manufacturer’s instructions from purified virus, infected FG cells and tissue samples. The E1 gene sequence of CsPaV and VP gene sequence of CsPV was used to design primers (Table 1) for virus detecting. Subsequently, the virus DNA samples were subjected to PCR (35 cycles of 95°C for 5 min, 94° C for 1 min, and 58°C for 1 min, and 72°C for 1 min, followed by extension at 72°C for 10 min), which produced a 654 bp and 1498 bp PCR product for CsPaV and CsPV, respectively.

Inverse PCR was performed to evaluate the circular genome of CsPaV. Primers used in inverse PCR are shown in Table 1. The virus DNA samples were subjected to PCR (35 cycles of 95 °C for 3 min, 94 °C for 10 s, and 48.2 °C for 10 s, and 72 °C for 1 min, followed by extension at 72°C for 7 min), which produced a 2046 bp PCR product.

### TaqMan probe-based Quantitative PCR

A TaqMan probe-based quantitative PCR was developed and used for detection and quantification of CsPaV and CsPV in infected Chinese tongue soles tissue samples and FG cells. Virus genomic DNA was extracted by DNA viral kit mentioned above. The primers and probes used in the study are shown in Table 1. A Premix Ex Taq™ (Probe qPCR) (TaKaRa, Japan) was used to perform qPCR following manufacturer’s instructions and each reaction consisted of 1 μl DNA template. The assay was performed at 95°C for 2 min, followed by 40 cycles of 95°C for 5 s and 60°C for 30 s. Each test was performed in triplicate together with a no template control in each run. To determine the copy numbers of the viruses, the standard curves for the TaqMan qPCR assays were generated using 10-fold dilutions of plasmid containing the DNA fragments of E1 gene from CsPaV and NS1 gene from CsPV. The virus copies were calculated according to the Ct values and the formula generated above.

### Prevalence of CsPaV and CsPV infection in Chinese tongue soles

To determine the prevalence of CsPaV and CsPV, a total of 123 Chinese tongue soles were collected from 9 locations in Tianjin, Hebei, Shandong province and Bohai Sea during 2021-2023 (Table 2). Livers, spleens and kidneys were sampled and subjected to quantitative PCR analysis to detect CsPaV and CsPV.

**Table 2.** The detecting results of clinical samples collected from different locations in China.

| Abbreviation of<br>sampling<br>location | Province<br>/location | Date of<br>collection | Temperature (°C) and<br>salinity (‰) | Size (cm) | Number of<br>fish | Positive<br>samples<br>of CsPaV | Positive<br>samples<br>of CsPV |
| --- | --- | --- | --- | --- | --- | --- | --- |
| HF | Tianjin | 15 Nov. 2021 | 17/20 | 37-40 | 29 | 29 | 16 |
| HS |  | 15 Oct. 2022 | 21/10 | 40-44 | 15 | 15 | 15 |
| LD |  | 3 Sep. 2023 | 22/18 | 24-27 | 10 | 10 | 10 |
| SY |  | 10 Nov. 2022 | 19/9 | 16-20 | 9 | 9 | 6 |
| YT |  | 11 May 2023 | 22/10 | 28-32 | 10 | 10 | 10 |
| LX |  | 18 May 2023 | 23/20 | 22-26 | 17 | 17 | 12 |
| TS | Hebei | 19 May 2023 | 21/13 | 17-19 | 10 | 10 | 10 |
| YT | Shandong | 21 May 2023 | 18/30 | 13-15 | 13 | 13 | 13 |
| Bohai Sea | Longitude<br>117.59-118.00 East,<br>latitude<br>39.10-39.08 North | 16 Oct. 2023 | 15/30 | 16-19 | 10 | 10 | 10 |

### Animal infecting experiments

Challenge experiments were performed using 96 healthy Chinese tongue soles. 48 fish in test group were challenged by intraperitoneal (IP) injection with 0.3 ml homogenate of diseased fish tissues prepared as described above and filtered through a 0.45 μm filter. 48 fish in the control group were injected IP with 0.3 ml Dulbecco’s PBS. All fish were held in tanks supplied with aerated water at 21°C and Salinity of 20‰ during the infection experiment. Clinical signs and mortality were checked daily.

## Results

### Diseases and clinical symptoms

Mortality occurred in farmed Chinese tongue soles in December 2021 in Tianjin China. Water temperature and salinity ranged from 17 °C to 20 °C and 16 ‰ to 20 ‰ at sites. Disease outbreak lasted for about 1 month (from December 16 2021 to January 16 2022) and the cumulative mortality reached 80.7% (Fig 1D). Clinical symptoms observed including swelling abdomen (Fig 1A-1), hemorrhages on the blind side of the fish (Fig 1A−2). The main internal signs included punctuate haemorrhages on the liver, swollen spleen and kidney filled with white nodules (Fig 1B-1, B-2).

**Fig 1.**
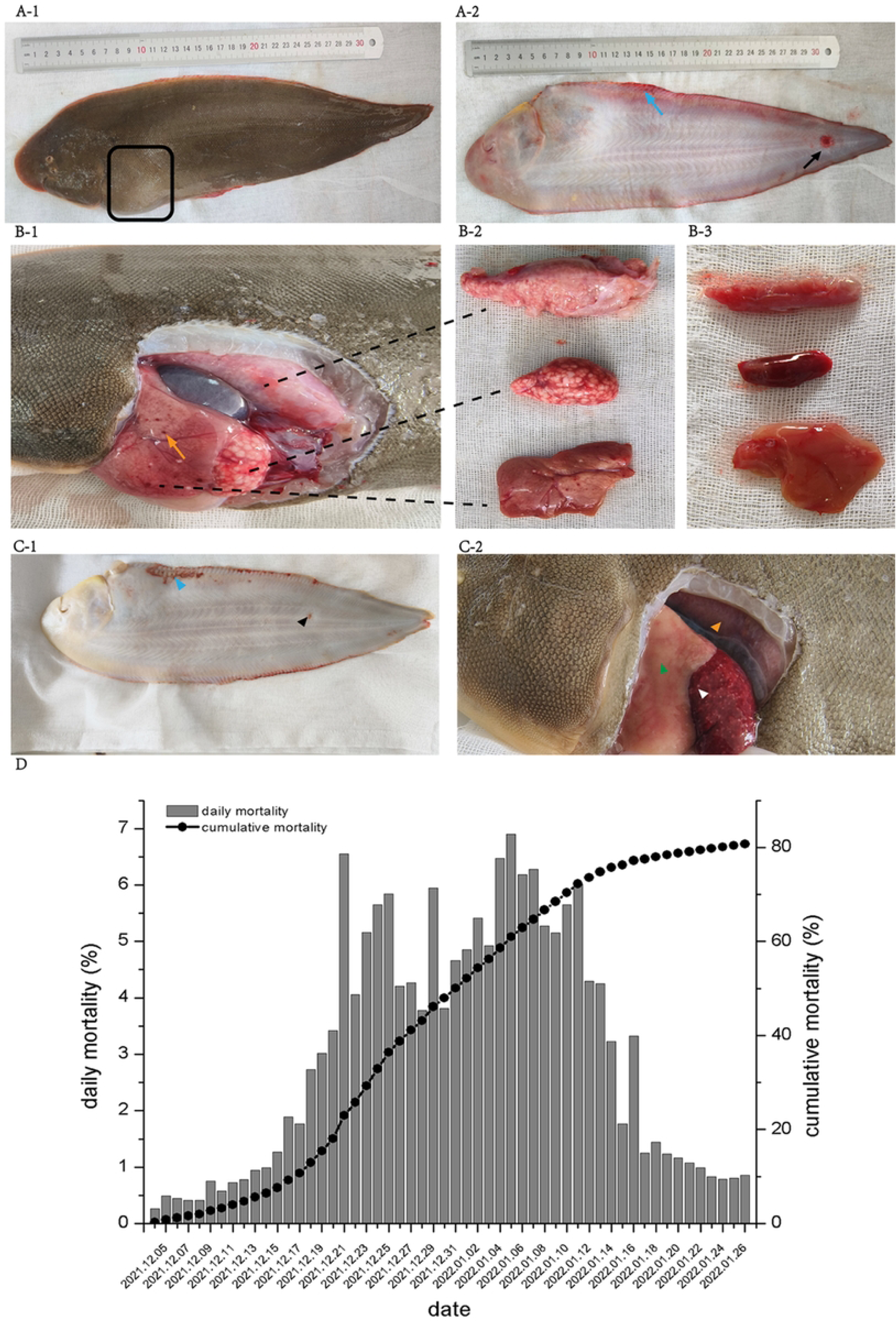
**Outbreaks and clinical symptoms of naturally diseased and experimentally infected Chinese tongue soles.** (A-1) Abdominal swelling (square frame) of the naturally diseased fish; (A-2) Hemorrhages near the tail (black arrow) and the fin bases (blue arrow) on the blind side of the fish; (B-1, B-2) Anatomical symptoms including swollen kidney and spleen filled with white nodules, punctate haemorrhages (orange arrow) on the liver; (B-3) kidney, spleen and liver from healthy Chinese tongue soles; (C-1) Hemorrhages (black arrowhead) and the fin bases (blue arrowhead) on the blind side of the experimentally infected fish; (C-2) Swollen spleen with white nodules (white arrowhead), slight swollen kidney (orange arrowhead), blood loss of liver (green arrowhead); (D) Statistics on daily and cumulative mortality.

**Fig 2.**
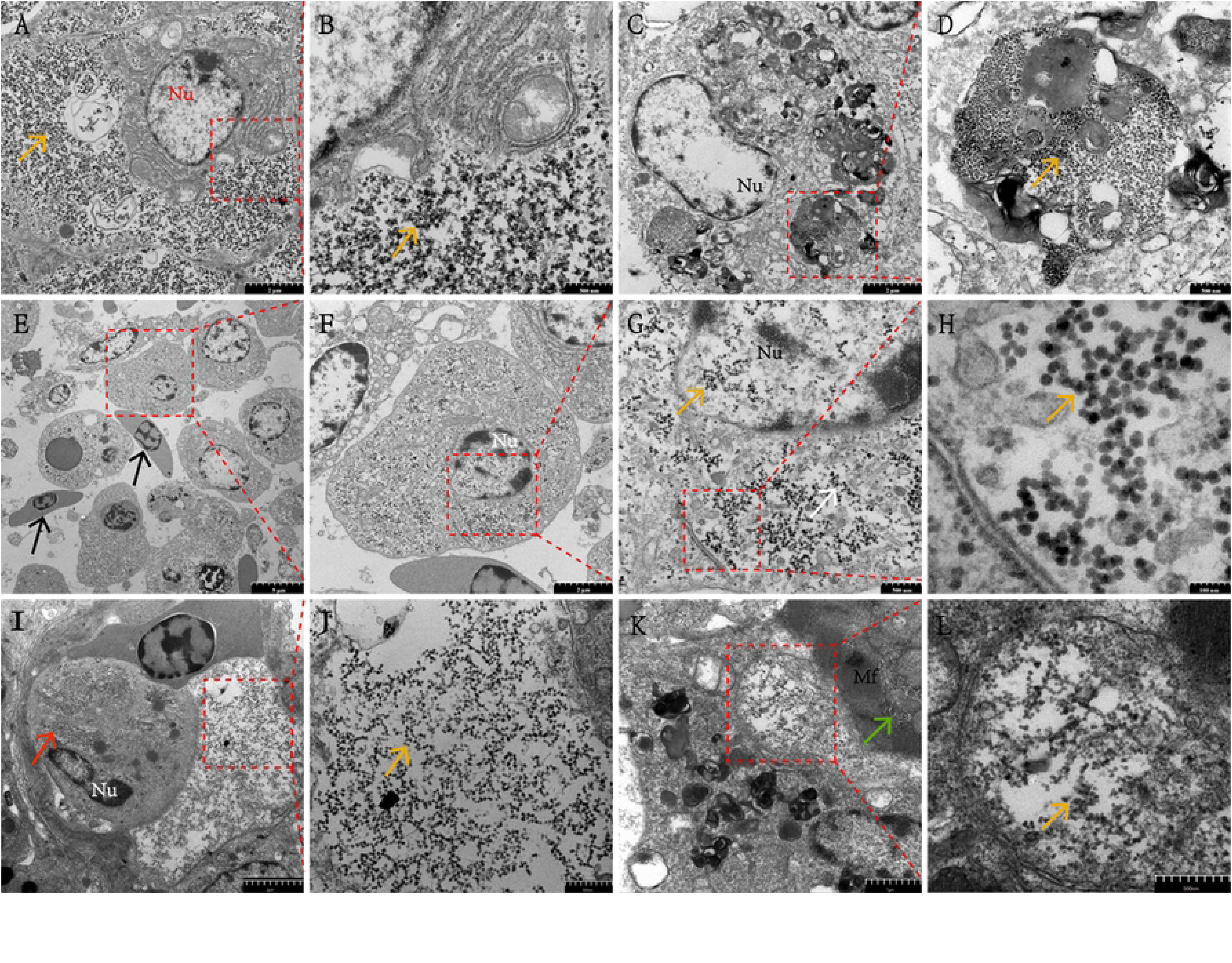
**Transmission electron micrographs of the viral particles from the tissues of naturally infected Chinese tongue soles.** (A, B) Liver. Large amount of virus particles were presented in the cytoplasm (orange arrow), Nu: nucleus (Bar, 2 μm) and high magnification of the red rectangular region in Panel A (Bar, 500 nm); (C, D) Kidney. Large, nearly circular inclusions in the cytoplasm near the nucleus (Bar, 2 μm) and the inclusion full of large number of virus particles (Bar, 500 nm); (E, F, G and H) Spleen. Bacterial pathogens (*vibrio*) (black arrow) located among the spleen cells and necrotic and lysed cells were observed (asterisk) (Bar, 5 μm), virus particles presented in cytoplasm (white arrow) and nucleus (orange arrow) (Panel F and G) (Bar, 2 μm and 500 nm), and high magnification of the virus particles with different size (about 28 nm in diameter showed with red arrow and 40 nm with green arrow) (Panel H, Bar, 100 nm); (I, J) Gill. Large numbers of virus particles are clustered in the cytoplasm (red arrow) or scattered in the interstitial spaces of cells (orange arrow) (Panel J, Bar, 500nm). (K, L) Heart. Virus particles occupied the interfibrillar spaces (orange arrow) and scattered in the cytoplasm (orange arrow) Mf: Myofibril (Bar, 1μm and 500 nm).

### Electron microscopy

Ultrastructurally, viral particles were observed from liver, kidney, spleen, gill and heart except brain tissues (Fig 2). Electron microscopy revealed that great number of spherical non-enveloped viruses scattered in the cytoplasm and nuclei of cells, interstitial and interfibrillar spaces. Moreover, large nearly circular inclusions filled with viral particles were observed in the cytoplasm near the nucleus (Fig 2C and D). Small amounts of co-infecting bacterial pathogens (*vibrios*) and lysed cells were also observed in the splenocyte (Fig 2E).

### Virus isolation and replication

Cell morphology change and CPE was examined at 3 days post-infection (dpi) (Fig 3A). Typical CPE including cell shrinkage, rounding and cytoplasmic vacuolization was observed at 5 dpi 2^nd^ cell culture passage (Fig 3B). The FG cells were rounding and floating in the medium at 8 dpi 3^rd^ cell culture passage (Fig 3C). No CPE was found in uninfected FG cells (Fig 3D). The TCID_50_ was determined as 10^-10.75^/mL at the 2^nd^ cell culture passage by 8 dpi.

**Fig 3.**
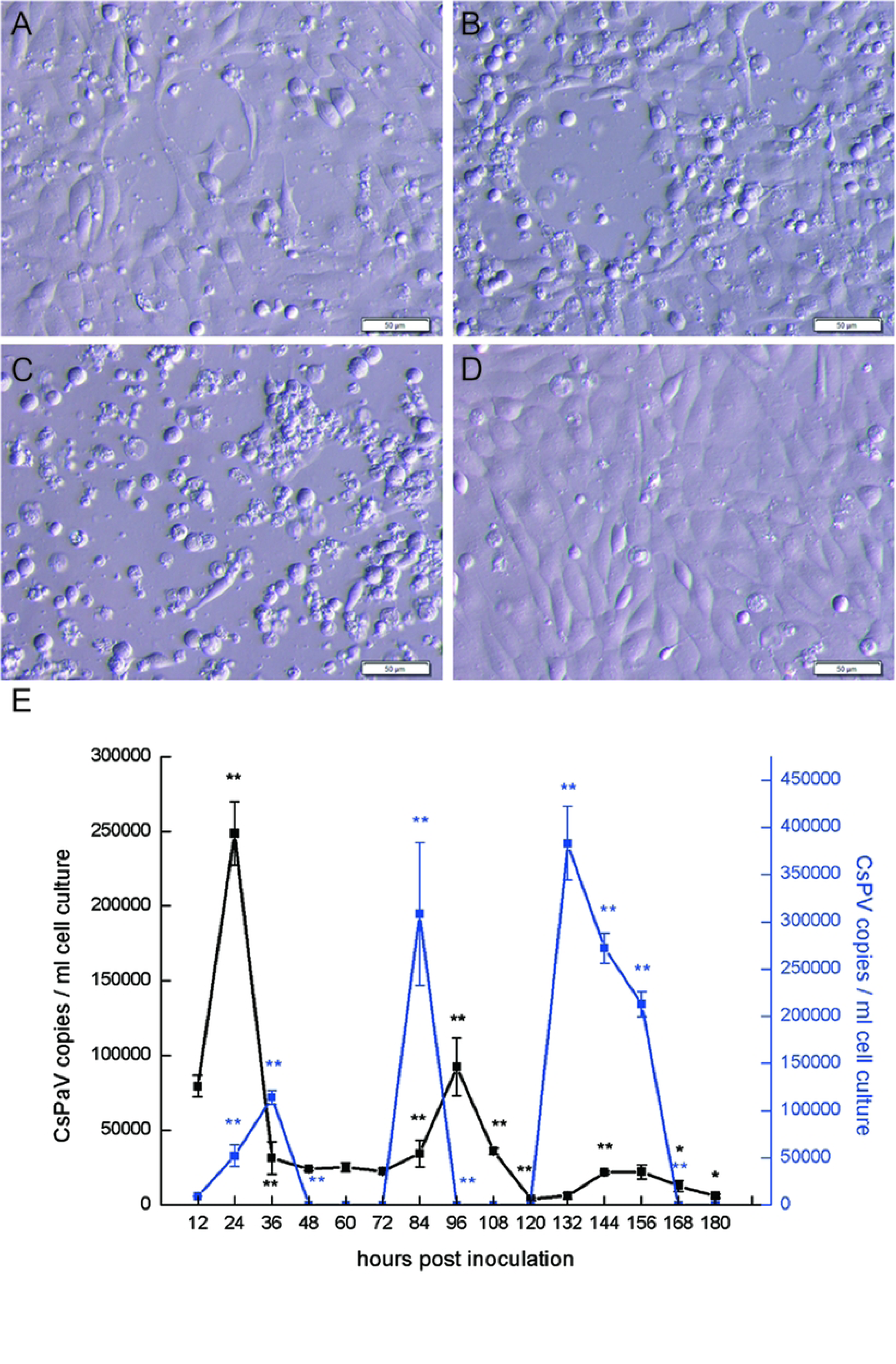
**Cytopathic effect (CPE) on Flounder gill cells (FG) induced by CsPaV and CsPV and replication dynamic of the two viruses on FG cells.** (A, B, C) FG cells infected with CsPaV and CsPV at passage 2, 3 days (A) and passage 2, 5 days (B) and passage 3, 7 days (C); (D) FG cells at passage 1, 3 days; (E) Replicating dynamics of CsPaV and CsPV in infected FG cells at different time point. The data represented means of three replicates of cell cultures. The data were analyzed by Student’s *t* test (*P*<0.05, one asterisks; *P*<0.01, two asterisks).

To determine the replication dynamics of CsPaV and CsPV in FG cells, viral copy numbers were analyzed using TaqMan probe-based Quantitative PCR. The results showed that both CsPaV and CsPV completed three replicating cycles within 180 hours (Fig 3E). The copy numbers of CsPaV and CsPV increased significantly at 24 hours post infection (hpi). CsPaV reached the first proliferation peak (10^5.39±0.037^/ml) at 24 hpi, while the first proliferation peak (10^5.06±0.028^/ml) of CsPV appeared at 36 hpi. Then, the viral copy numbers was decreased significantly to 10^4.48±0.15^/ml for CsPaV and undetectable for CsPV in 12 hours. The second proliferation peak of CsPV occurred at 84 hpi with concentration 10^5.48±0.11^/ml and decreased to an undetectable degree at 96 hpi. While the copy number of CsPaV increased to 10^4.96±0.091^/ml at 96 hpi and decreased to 10^3.62±0.086^/ml at 120 hpi. And then, the viral copy numbers of CsPV began to increase significantly after 24 h of silencing, reaching the third proliferation peak (10^5.58±0.04^/ml) at 132 hpi, and decreased significantly again to an undetectable degree at 168 hpi. While CsPaV copies reached a relative small replication peak (10^4.34±0.018^/ml) at 144hpi and decreased to 10^3.79±0.09^/ml at 180 hpi.

### Histopathology

Liver, spleen, kidney and intestine tissues from experimentally infected fish for 3, 5 and 7 days were used to analyze the pathological changes caused by viral infections. In liver, marginated and fragmentized nuclear, some necrotic and vacuolated hepatocytes accompanying hepatic sinusoidal without erythrocytes were observed at 3 dpi. Numerous necrotic, vacuolated hepatocytes and enlarged empty hepatic sinusoidal presented at 5 dpi, and then lysed hepatocytes, aggregated nucleus and necrotic white pulp appeared at 7 dpi. The hepatocytes were typical in arrangement and stain evenly and the hepatic sinusoids contained a large number of erythrocytes in un-infected liver (Fig 4A). In spleen, ruptured blood vessels and scattered blood cells, lysed and vacuolated splenocytes occurred at 3 dpi. Necrotic splenic white pulp, virus inclusions with a decrease in the number of splenocytes were found at 3 and 5 dpi. The splenocytes were stained evenly with regular cell shape in healthy spleen (Fig 4B). In kidney, numerous necrotic and vacuolated kidney cells, edematous renal tubules were observed at 3 dpi which showed an acute infection. Atrophied, vacuolized renal tubules, necrotic white pulp and some virus inclusions accompanied a decrease in the number of kidney cells were found at 5 and 7 dpi. The cells stained evenly and the renal tubules structurally intact in the healthy kidney (Fig 4C). In the infected intestine, intestinal mucosa underwent a progression from edema, erosion to ulceration during the virus infection. The atrophic, disarranged and denatured and exfoliated epithelial cell were observed at different periods of infection, inflammatory cell infiltration were present during the virus infection. The intestinal mucosa was intact, epithelial cells were well arranged and stained evenly in healthy intestine (Fig 4D).

**Fig 4.**
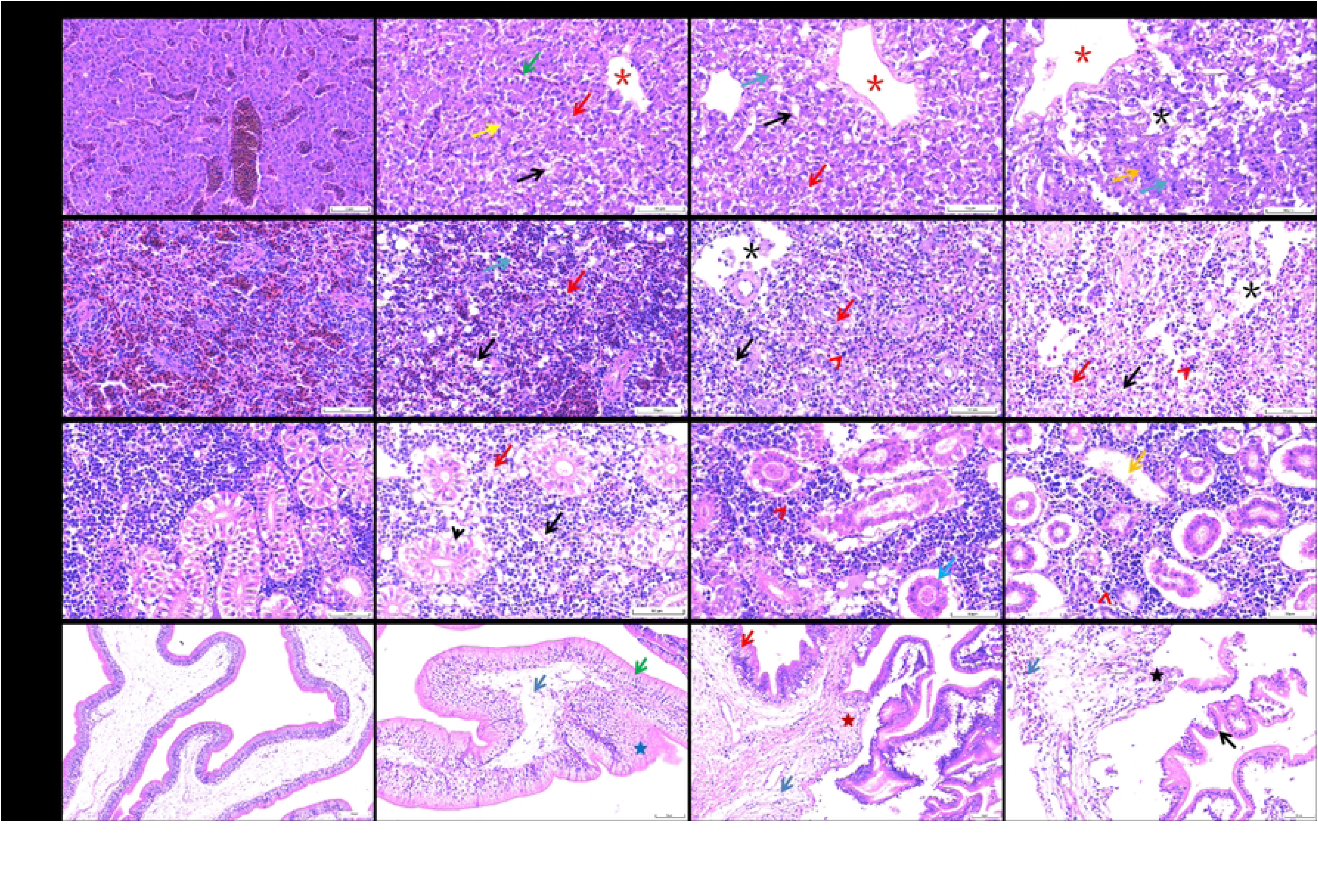
**Pathological analysis.** (A) Liver; marginated (green arrow) and fragmentized (yellow arrow) nuclear, necrotic (red arrow) and vacuolated hepatocytes (black arrow); enlarged empty hepatic sinusoidal (red asterisk); lysed (blue arrow) hepatocytes, aggregated nucleus (orange arrow) and necrotic white pulp (black asterisk) presented in the infected liver; (B) Spleen; large amounts of lysed (red arrow) and vacuolated splenocytes (black arrow), ruptured blood vessels and scattered blood cells (blue arrow) presented in the spleen at 3 dpi; more lysed and vacuolated splenocytes, virus inclusions (red arrowhead) and necrotic splenic white pulp (black asterisk) accompanied by a decrease in the number of splenocytes were found at 5 and 7 dpi. (C) Kidney; necrotic (red arrow) and vacuolated (black arrow) kidney cells and edematous renal tubules (black arrowhead) accompanied a decrease in the number of kidney cells were observed at 3 dpi; atrophied (blue arrow) and vacuolized (orange arrow) renal tubules and some virus inclusions (red arrowhead) were found in the kidney at 5 and 7dpi; (D) Intestine. edematous (blue pentagram), eroded (red pentagram) and ulcerated (black pentagram) mucosa, atrophic (green arrow), disarranged (red arrow), denatured and exfoliated (black arrow) epithelial cell; were observed at 3, 5 and 7 dpi and inflammatory cell infiltration (blue arrow) were present during the virus infection.

### Genome sequencing analysis

SISPA-PCR combined with high-throughput sequence approach (Fig 5A) was used to analyze the full viral genome sequence. No results were obtained from the RNA template. Unexpectively, two kinds of virus-associated proteins belonging to the *Papillomaviridae* and Parvovidae families respectively were found when comparing the filtered sequences with other viral proteins in the GenBank database using BLASTX. And then two complete genomes named *Cynoglossus semilaevis* papillomavirus (CsPaV) and *Cynoglossus semilaevis* parvovirus (CsPV) were assembled using Soap *denovo* 2.0, and the sequences of the genomes were confirmed by overlap PCR or inverse PCR followed by Sanger sequencing.

**Fig 5.**
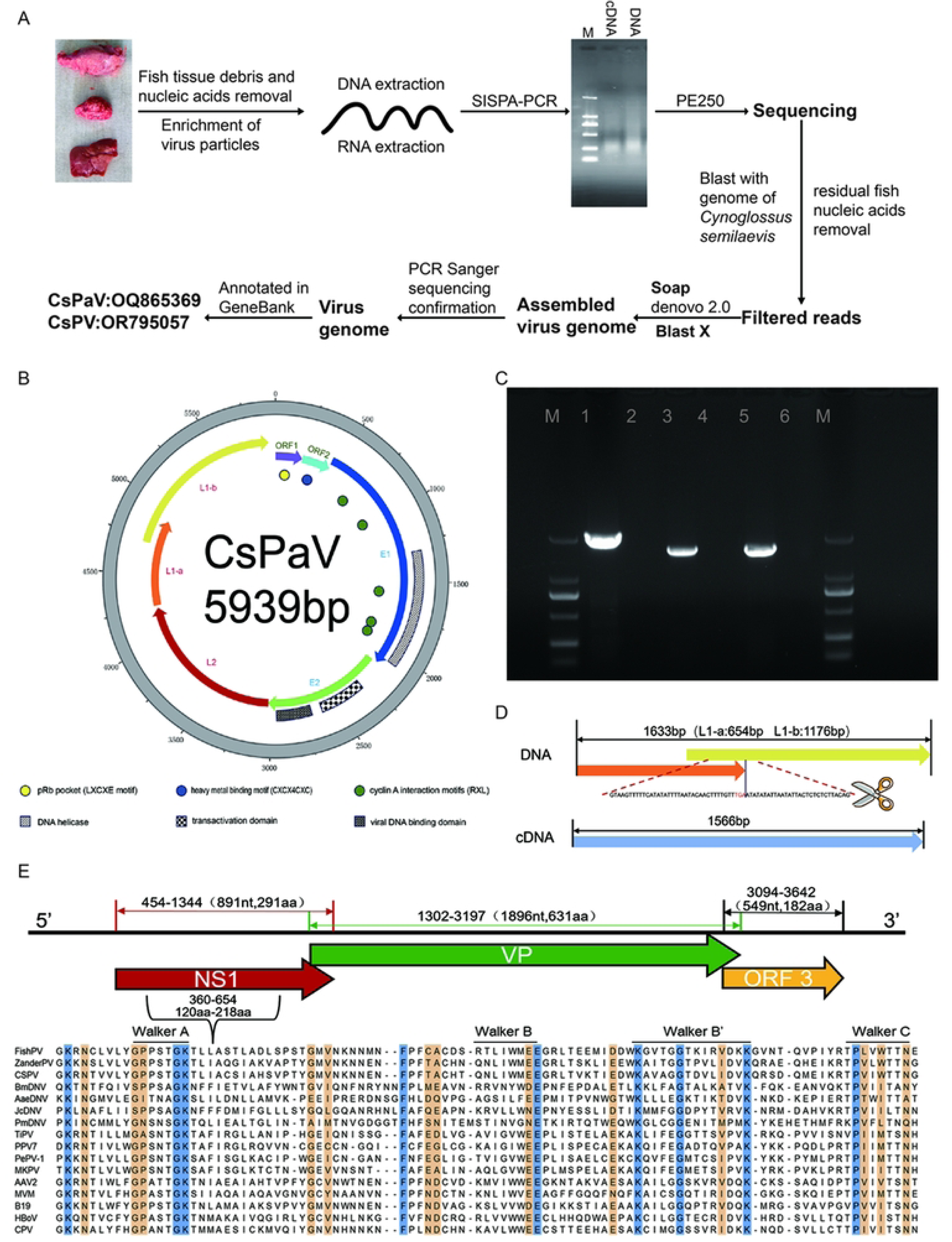
**Identification and genomic analysis of the causative viruses of diseased Chinese tongue soles.** (A) Workflow to identify the genomes of CsPaV and CsPV from the infected fish; (B) Circular graphs of the genome of CsPaV. Outer scales are numbered in kilobase pairs in a clockwise direction. The predicted E1, E2, L2 and L1 genes are showed in different colors by arrows; (C) Inverse PCR on extracted genomic DNA of CsPaV infected (lane 1) and uninfected (lane 2) liver tissues of fish, RT-PCR on RNA extracted from CsPaV infected (lane 3) and uninfected (lane 4) FG cells, PCR on genomic DNA extracted from CsPaV infected (lane 5) and uninfected (lane 6) FG cells; (D) Diagram showing CsPaV-L1 containing two ORFs (L1a and L1b), with the spliced sequence (67 bp) indicated; (E) Genome organization of CsPV (The NS1, VP and ORF3 are shown in different colors) and alignment of conservative helicase domains including Walker A, B, B^’^ and C box of NS1 protein.

The complete genome of CsPaV was 5939 bp and had a GC content of 36.95%. The results of the inverse PCR revealed that CsPaV had a circular genome and it was not an endogenous viral element in the genome of the host *Cynoglossus semilaevis* (Fig 5C). The genome contained four predicted protein-coding regions, which were typical and backbone proteins of papillomavirus (E1, E2, L1 and L2), lacking any of the oncogenes (E5, E6 and E7) [9] (Fig 5B). The E1 protein had a DNA helicase (aa 314-561) at C terminal [10] and five cyclin A interaction motifs (RXL; aa 90-92, aa 186-188, aa 383-385, aa 469-471, aa 477-479). While the E2 protein contained a transactivation domain (aa 84-171) and a viral DNA binding domain (aa 213-288) identified from other papillomavirus [11]. Moreover, two small open reading frames (sORF1 and sORF2) which showed no similarities to other proteins in the database were found at the region between the end of the L1 ORF and the beginning of the E1 ORF. The sORF 1 encoded 70 amino acids and had a pRb pocket (LXCXE motif). While the sORF2 ecoded 71 amino acids and had a heavy metal binding motif (CXCX_4_CXC). PDZ-binding motif which was correlated with oncogenic potential [12] was absent in both sORF1 and sORF2. Additionally, an E1 binding site (E1BS; ATNGTTN_3_AAGNAT) was found at non-coding area before sORF1 (nt 5498-5512). Interestingly, the encoding region of CsPaV L1 was composed of two ORFs (L1a and L1b), with L1a overlapping by 195 nt with L1b, when predicted by ORF finder (Fig 5B). The unusual structure suggested that the L1 protein might be expressed by spliced mRNA. To confirm it, the splicing sites were predicted by Net-Gene2 Server. The results showed that donor and acceptor splice sites were found at nt 617 (confidence 0.83) and nt 683 (confidence 0.97), respectively. To verify the prediction, the total RNA of CsPaV-infected tissues, FG cells and transfected FG cells were extracted, and then RT-PCR was conducted to amplify the L1 gene. The PCR products were sequenced and BLAST with the sequence obtained from the DNA template. The results showed that a 67 bp segment with high AT content (81.6%) was spliced in the cDNA template, and the splicing sites were at nt 617 and nt 683 as expected (Fig 5D). The complete genome sequence of CsPaV was deposited in GenBank (accession no. OQ865369).

A novel parvovirus (CsPV) was also found in the diseased fish. The nearly complete genome was 3663 bp and contained three ORFs, encoding the well-known proteins of NS1 and VP of parvovirus, and an unknown ORF3 respectively. The VP gene overlapped by 32 nt with the NS1 gene, the unknown ORF3 overlaped by 103 nt with the VP gene (Fig 5E). The NS1 was predicted to be 891 bp (296 aa) in length which was significantly shorter than other parvovirus in subfamily *Parvoviridae* (569-651 aa). It contained a helicase domain (aa 120-218) including the conserved ATP- or GTP-binding Walker loop motifs, Walker A loop aa motif (GxxxxGKT/S; 127GPPSTGKT137), and the Walker B (xxxxEE; 168LIWMEE175), Walker B’ (KxxxxGxxxxxxxK; 185KAVAGGTDVLIDVK200), and Walker C (PIxIXXN; 210PVIWTTN218) aa motifs [13]. The HuH motif which were widely present in the NS1 was absent in CsPV [14]. The VP protein was 631 aa in length which was similar with other members in subfamily *Parvovirinae* (537-781 aa). Similar with other identified fish parvovirus [15, 16], the phospholipase A2 motif which were widely present in the VP protein of many parvoviruses [17] was not found in CsPV. Besides, an unknown ORF3 encoding 182 aa was found and showed no significant matches with other parvoviruses by BLASTP. The genome sequence of CsPV was deposited in GenBank (accession no. OR795057). The sequencing reads were deposited into the Sequence Read Archive (SRA) under accession number PRJNA1040012.

### Phylogenetic analysis

A phylogenic tree was constructed basing on the nucleotide sequence of the L1 gene (http://pave.niaid.nih.gov) (S1 Fig). It showed that CsPaV clustered with other fish papillomavirus, while formed a separate branch with genus *Nunpapillomavirus* of subfamily *Secondpapillomavirinae* and clustered with other unclassified fish papillomavirus. Besides, phylogenetic analysis based on the amino acid sequence of the L1 protein (Fig 6 A) revealed that the L1 protein of CsPaV shared the highest similarity of 53.45% with *Papillomaviridae* sp. Haddock_c6033 (GenBank accession no. : QYW06024.1). According to the genus demarcation Criteria of the International Committee on Taxonomy of Virus (ICTV) for family *Papillomaviridae* (viruses with >70% identity in the L1 sequence belong to the same species; those >60% identity belong to the same genus), we proposed that CsPaV belonged to a new genus of this group. As for CsPV, a phylogenetic tree was constructed based on the NS1 protein amino acid sequences of 20 representative parvovirus in the subfamily *Parvovirinae* and Hama*Parvovirinae* included in the family *Parvoviridae* [Fig 6B]. It revealed that CsPV clustered with *Parvovirinae* sp. isolate fi102par1 (GenBank accession no. : OP36693) and zander/M5/2015/HUN, sharing 51.15%, 34.85% similarity, respectively. Based on ICTV *Parvoviridae* Study Group, all parvoviruses in a genus should be monophyletic and encode NS1 proteins that >30% identical to each other at the amino acid sequence level [18], we speculated that CsPV belonged to a new genus with *Parvovirinae* sp. isolate fi102par1 and zander/M5/2015/HUN in subfamily *Parvovirinae*.

**Fig 6.**
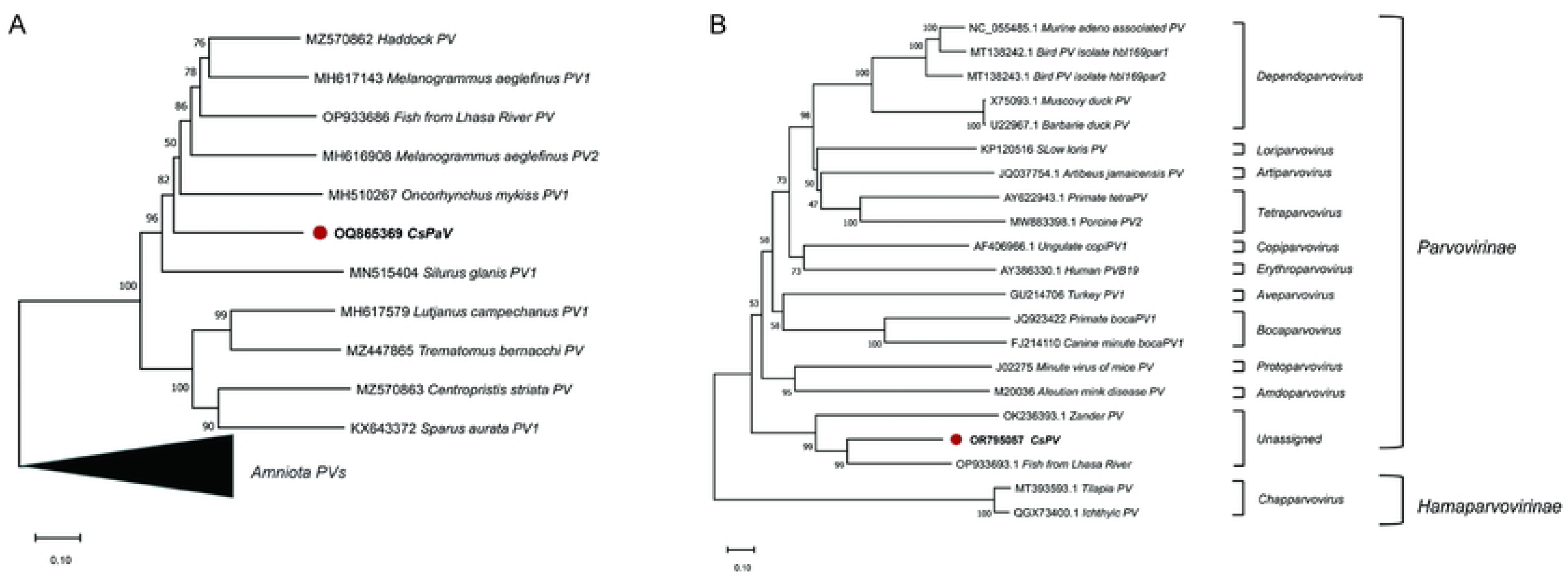
**Phylogenetic analysis of CsPaV and CsPV.** (A) Phylogenetic tree of papillomaviruses based on aligned amino acid sequences of the CsPaV L1 protein and the L1 proteins of other fish papillomaviurses; (B) Phylogenetic tree of parvovirus based on amino acid sequence alignment of CsPV NS1 protein and the NS1 proteins of other representatives of 10 genera in the subfamily *Parvovirinae* and 2 unassigned fish parvovirus in subfamily *HamaParvovirinae*. The phylogenetic trees were constructed using the Maximum Likelihood method with 100 bootstrap replicates. The scale bar means the genetic distance, number of substitutions per site.

### FISH detection

The mRNA expression of CsPaV and CsPV in the infected fish was analyzed by Paraffin-SweAMI-double FISH method and shown in Fig 7. The transcripts of the two viruses located mostly in cytoplasm (red / green) and a little in nucleus (purple /cyan). Moreover, CsPaV and CsPV could be detected in cytoplasm (yellow) and nucleus (white) of one cell.

**Fig 7.**
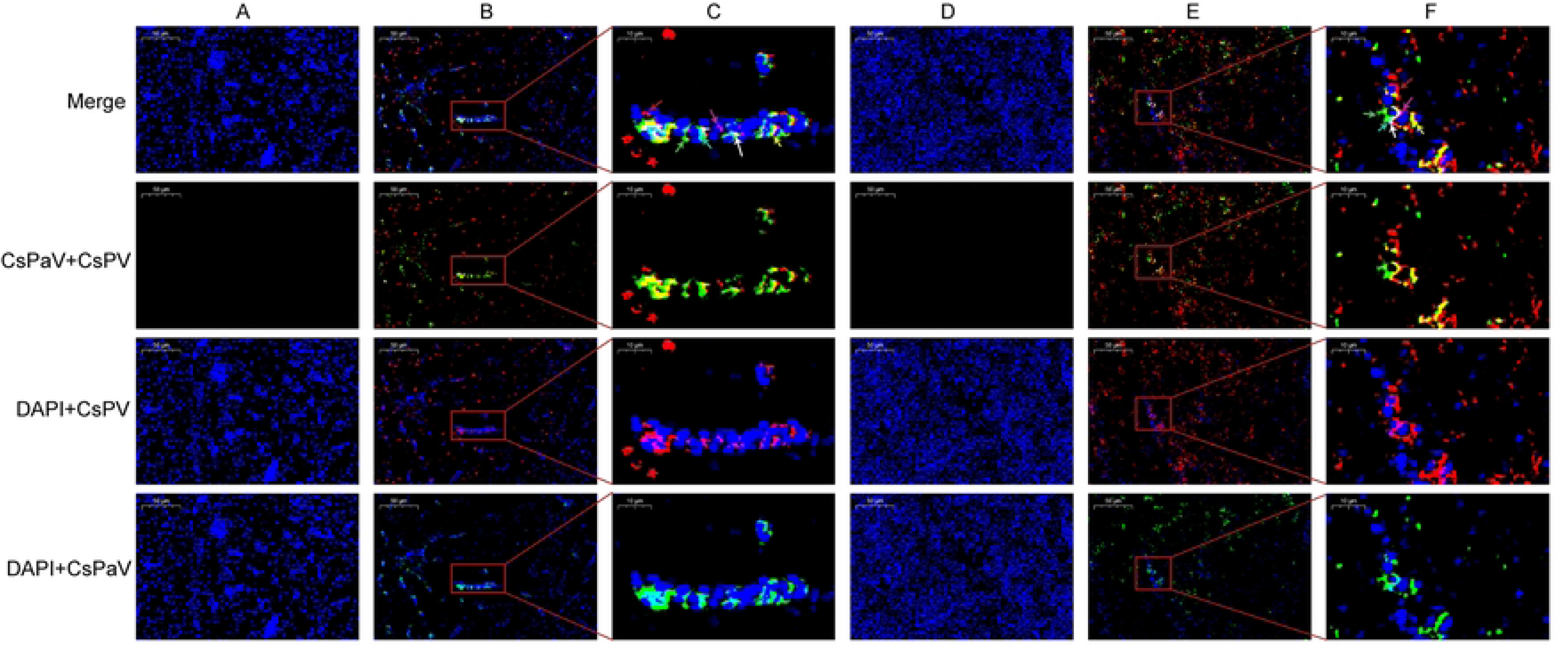
**Detection of the mRNAs of CsPaV and CsPV in infected liver and spleen of Chinese tongue soles by FISH.** (A) Uninfected liver (Bar, 50 μm); (B) Infected liver (Bar, 50 μm); (C) High magnification of the red rectangular region (Bar, 10 μm); (D) Uninfected spleen (Bar, 50 μm); (E) Infected spleen (Bar, 50 μm); (F) High magnification of the red rectangular region (Bar, 10 μm). Tissue sections display red fluorescence for CsPV (Cy3-conjugated probes), green for CsPaV (FAM-conjugated probes) and blue for cell nucleus (DAPI). The mRNAs of CsPV and CsPaV was detected in cytoplasm which showed red (red arrow) and green (green arrow) and nucleus showed purple (purple arrow) and cyan (cyan arrow). The mRNAs of the two viruses co located in the cytoplasm showed yellow (yellow arrow) and nucleus showed white (white arrow) of one cell.

### Viral loading and replicating dynamics in different tissues

TaqMan probe-based quantitative PCR of CsPaV and CsPV was performed to analyze viral loading and replicating dynamics in nine tissues (liver, spleen, kidney, intestines, gill, muscle, heart, mucus and eye) sampled at 1,2,3,5 and 7 dpi. The results showed that the concentration of the two viruses reached maximum values by 5 or 7 dpi in most examined tissues and the virus copy amounts of CsPaV were much higher than that of CsPV (Fig 8E and S2 Fig). In liver, CsPaV viral copies decreased from 10^6.22±0.023^ / mg by 1 dpi to 10^5.66±0.054^ / mg by 2 dpi and increased significantly to 10^5.84±0.042^ / mg by 3 dpi and 10^6.73±0.04^ /mg by 5 dpi, and then decreased to 10^6.38±0.085^ / mg by 7 dpi (Fig 8E). The similar replicating dynamics were observed in muscle, intestine, heart and gill (S2 Fig). While in kidney, the viral copies increased slightly from 10^6.68±0.083^ / mg by 1 dpi to 10^6.86±0.011^ /mg by 2 dpi, decreased to 10^6.26±0.03^ / mg by 3 dpi, and then increased significantly to 10^7.39±0.041^ /mg by 7 dpi. It indicated that CsPaV completed two replicating processes within 7 days and the releasing viral amounts at the first peak (by 2 dpi) were much lower than that of the second peak (by 5 or 7 dpi). The same viral proliferations were observed in mucus and eye (S2 Fig). As for CsPV, the viral copies decreased from 10^2.59±0.21^ /mg by 1 dpi to undetectable by 3 dpi, and then increased significantly to 10^3.52±0.27^ / mg by 3 dpi and decreased to 10^2.60±0.21^ / mg by 7 dpi in the liver. The same dynamics was observed in heart. In kidney, the virus increased to a peak with 10^3.43±0.14^ / mg by 3 dpi and decreased to 10^3.10±0.12^ / mg by 7 dpi (Fig 8E). However, a different virus replicating curve was observed in spleen which showed that the virus copies kept a higher concentration at all the sampled points with a maximum value 10^3.37±0.31^ / mg by 5 dpi and then decreased to 10^3.32±0.01^ / mg. More interestingly, the CsPV remained undetectable by 1, 2 and 3 dpi and increased significantly with a maximum value by 5 dpi and decreased to nearly undetectable again by 7 dpi in intestine, muscle, gill, mucus and eye (S2 Fig).

**Fig 8.**
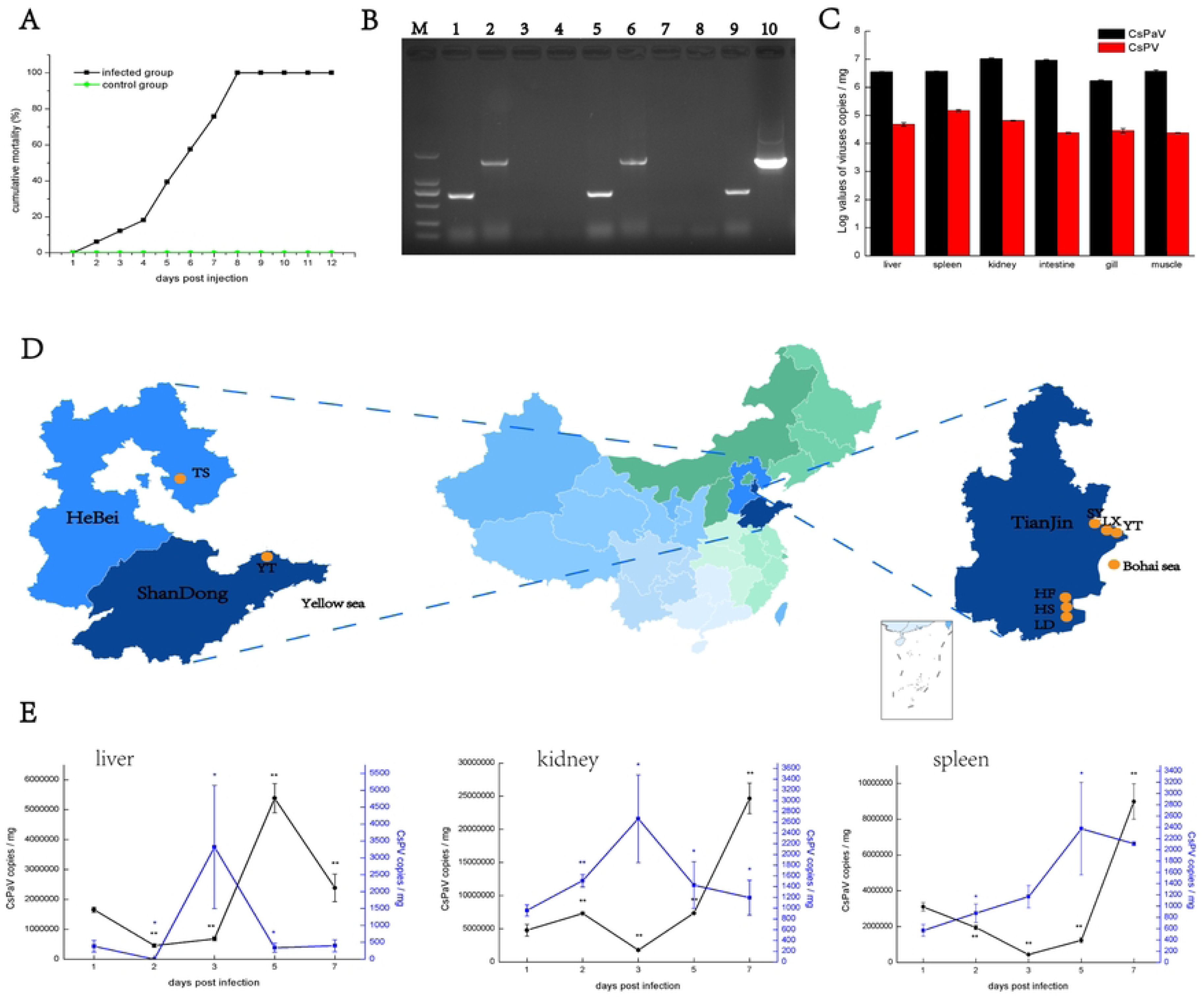
**Distribution and replicating dynamics of CsPaV and CsPV in different tissues and cumulative mortality of artificially infected fish and prevalence of the two viruses infection in different locations of China.** (A) Cumulative mortality of Chinese tongue soles infected with CsPaV and CsPV; (B) PCR detection of CsPaV and CsPV in the experimental group and control group. Lane M: DL2000 Marker; Lane 1 and 2: CsPaV and CsPV in kidney sample of experimental group; Lane 3 and 4: CsPaV and CsPV in kidney sample of control group; Lane 5 and 6: negative control; Lane 7 and 8: positive control of CsPaV and CsPV; (C) Tissue distribution loading of CsPaV and CsPV in naturally infected Chinese tongue soles. The data represented means of three fish samples, and the error bars indicated standard deviations. (D) Locations of diseased Chinese tongue soles collected from Tianjin, Shandong, Hebei province and Bohai sea. Yellow circles indicated samples that were positive for CsPaV and CsPV, and green indicated samples that were positive for CsPaV. (E) CaPaV and CsPV genome copies in liver, kidney and spleen tissues of experimentally infected fish at different time point (1, 2, 3, 5 and 7 dpi). The data represented means of three fish samples. The data were analyzed by Student’s *t* test (*P*<0.05, one asterisks; *P*<0.01, two asterisks).

### Distribution and prevalence of CsPaV and CsPV

The liver, spleen, kidney, intestine, gill and muscle of naturally infected Chinese tongue soles were collected and virus loading were analyzed by probe-based quantitative PCR. The results showed that both viruses had high loads in the examined tissues. CsPaV had the highest loads in kidney (10^7.02±0.02^ / mg) followed by intestine (10^6.96±0.03^ / mg), and almost equal loads were observed in liver spleen (10^6.57±0.01^ / mg), (10^6.55±0.01^ / mg) and muscle (10^6.57±0.05^ / mg), with gill (10^6.23±0.03^ / mg) having the least amount of virus. For CsPV, the loads in the tissues were significantly lower than those of CsPaV. The virus copies of CsPV were much higher in spleen (10^5.17±0.04^ / mg), kidney (10^4.81±0.01^ / mg) and liver (10^4.68±0.06^ / mg) than in gill (10^4.45±0.08^ / mg), intestine (10^4.37±0.02^ / mg) and muscle (10^4.37±0.01^ / mg) (Fig 8C).

Total of 123 samples from 9 fish farms in Tianjin, Hebei and Shandong provinces, and wild Chinese tongue soles from Bohai Sea (E 117.59-118.00, N 39.10-39.08) were collected, and then the two viruses were detected using probe-based quantitative PCR. The results showed that all the samples from the 9 locations were positive for CsPaV, and the positive rate of CsPV ranged from 55%-100% with various virus loads, and the two viruses cocurrented in 82.9% sampled fish (Table 2)(Fig 8D).

### Animal experiments

Homogenate of diseased fish tissues was used to intraperitoneally inject the healthy Chinese tongue soles with with 10^7.71^ copies for CsPaV and 10^6.16^ copies for CsPV per fish determined by probe-based quantitative PCR. The fish showed feeding disorders, and decreased homeostasis by 4-6 dpi. Clinical symptoms such as hemorrhagic spots on the body surface (Fig 1C-1), swollen spleen with white nodules in spleen, swollen kidney and liver losing blood (Fig 1C-2) were observed in the infected fish. The cumulative mortality reached 100% at 8 dpi (Fig 8A). All the fish in the control group remained asymptomatic. Two fish were selected randomly from the experimental and control group respectively at 5 dpi to detect the CsPaV and CsPV using PCR method. The results showed that samples in the experimental group were PCR positive for CsPaV and CsPV, while samples from the control group were negative (Fig 8B).

## Discussion

Chinese tongue sole is an economically important fish for marine aquaculture, and in recent years, the outbreak and prevalence of an emerging viral disease has severely limited the healthy and sustainable development of the fish aquaculture industry. In fact, as early as 2015, viral particles was observed in the tissues of Chinese tongue sole suffering from splenic necrosis disease [5], but the viral pathogen has not been identified. Although some novel viral agents have been characterized by viral metagenomics [19–20], the technique is frequently hampered by the relatively lower virus load and higher background nucleic acids in the tissues of the host. More importantly, the genomes of viruses with single-stranded nucleic acids cannot be analyzed by this method because the single stranded nucleic acid cannot be processed directly to small fragments to construct the sequencing library. Improved technologies including SISPA and virus discovery cDNA-AFLP (VIDISCA) facilitated the discovery of several novel agents [20–22]. However, the huge sequence background of the host is still a big obstacle for novel virus identification. In the present study, we proposed a new strategy to analyze the genome of novel virus which the key procedure was filtering and removing the host nuclear acids by blasting the sequences produced from the SISPA products with the genome of *Cynoglossus semilaevis* [8]. It was proved that the approach could rapidly and accurately identify novel viruses whose host genomes have been sequenced.

Novel parvovirus (CsPV) and papillomavirus (CsPaV) were proved to be the pathogenic pathogens causing massive mortality of Chinese tongue soles. Parvovirus is small, resilient, non-enveloped viruses with linear, single-stranded DNA [36]. Phylogenetic analysis showed that CsPV belonged to a new genus in subfamily *Parvovirinae* of family *Parvoviridae*. To date, a total of five fish parvoviruses, including CsPV, have been identified, while parvoviruse from Lhasa river (GeneBank accession no. OP933693.1) was identified by megagenomics and no other information was supplied. Zander PV [15] and ichthyic PV [16] were obtained via fecal collection. So it remains unclear whether the three viruses are disease-causing agents. TiPV belonging to genus *Chapparvovirus*, subfamily Hama*Parvovirinae* was proved to be the causative agent of tilapia [14]. We suggested that CsPV belonging to a new genus of subfamily *Parvovirinae* was the first disease-causing parvovirus identified from marine cultured flatfish, suggesting evolutionary adaptations to high salinity of CsPV and broadening the presence range of parvovirus.

Another novel disease-causing virus was CsPaV. Papillomaviruses are small icosahedral, non-enveloped viruses with circular, double stranded DNA [30]. Phylogenetic analysis revealed that CsPaV clustered together with other identified fish papillomavirus, suggesting a possible monophyletic origin of the group. CsPaV belonged to a new genus in subfamily *Secondpapillomavirinae* of family *Papillomaviridae*. So far, papillomaviruses have been found in eight fish species including gilthead sea bream (*Sparus autata*) [23], wels catfish (*Silurus glanis*) [24], black sea bass (*Centropristis striata*), emerald notothen (*Trematomus bernacchii*), haddock (*Melanogrammus aeglefinus*) [25], rainbow trout (*Oncorhynchus mykiss*), red snapper (*Lutjanus campechanus*) [26] and fish from Lhasa river (GeneBank accession no. OP933686.1). Most of the determined fish papillomaviruses have been obtained by megagenomic sequencing, and it remains unclear whether they are disease-causing agents of the fish. While Sparus aurata papillomavirus 1 (SaPV1) and Silurus glanis papillomavirus 1 (SgPV1) were reported to be one of the agents caused typical signs of diseases in fish such as papillomalike lesions or papilloma-like epidermal hyperplasia on the skin, which were consistent with the typical syndromes from papillomavirus infection due to they are epitheliotropic viruses inducing infections in the stratified squamous epithelia of skin and mucosal membrane [27]. However, we observed “tumor like” signs and high CsPaV loads in the kidney and spleen tissues of diseased Chinese tongue soles. Some research revealed that papillomavirus could bind to heparin sulfate proteoglycans on either the epithelial cell surface or basement membrane through interactions with the L1 protein [28,29] and its replication cycle is tightly coupled to the differentiation state in infected squamous epithelia [31]. We speculated that CsPaV might have evolved new strategies to bind and entry the cells and then replicating in the cells of visceral tissues due to the altered histophilicity.

Viral co-infections are common in nature. However, studies and available data on viral co-infections in fish aquaculture are limited [42,43]. In recent years, infectious events of fish papillomavirus co-infected with other viruses were found in fish aquaculture, such as SaPV1 co-infected with iridovirus and polyomavirus [23], SgPV1 co-infected with herpesvirus [24]. And CsPaV co-infected with CsPV in the present study. There are five viral interaction patterns including interference, synergy, noninterference, dependence assistance and host–parasite relation. Currently, we cannot elucidate the co-infecting pattern of CsPV and CsPaV because we did not obtain enough single virus infected samples. Epidemiological investigation revealed that CsPaV and CsPV cocurrented in most sampled fish (82.9%), suggesting co-operative effects on the emerging disease. Moreover, the mRNAs of CsPaV and CsPV could be detected in one cell by FISH, hinting at the possibility of two viruses interacting each other. Viral replicating dynamics showed that CsPaV and CsPV both completed at least one replicating cycle in infected FG cells or tissues. The copy numbers of CsPV are much higher than that of CsPaV in the infected FG cells, while the opposite effect appears in the infected tissues, making it complicate to address the interacting pattern of the two viurses. It was reported that adeno-associated virus (AAV) which is a kind of parvovirus has shown to interact with human papillomavirus (HPV), and it could reduce the risk of developing HPV-16-induced cervical cancer [32–33]. While HPV16 were shown to play a significant role in enhancing AAV replication, and AAV replication could interfere with the replication and oncogenic potential of papillomavirus [34–35]. Whether CsPaV and CsPV have similar interacting pattern is unknown, and the mechanism requires further research.

Unusual genome organizations were found in CsPV and CsPaV. There are four typical genome structures contained two major ORFs encoding NS1 and VP in family *Parvoviridae* (https://ictv.global/report/chapter/Parvoviridae/Parvoviridae). While the genome of CsPV contains, in addition to the ORFs of NS1 and VP, an unknown ORF3, encoding 182 amino acids at the 3’ end, with no homology to any nucleotide or amino acid in GenBank. Moreover, the NS1 gene and the VP gene, the VP gene and the unknown ORF3 have a short overlap each other, which is uncommon in subfamily *Parvovirinae*. In a recent research, an unknown 125-aa-long protein at 5’ end was also found in a parvovirus (zander/M5/2015/HUN) identified from faecal specimens of zander, and the NS1 also overlapped with VP by 361 nt, while VP and the unknown protein was separated by 42 nt [16]. Moreover, a parvovirus (*Parvovirinae* sp. isolate fi102par1, GenBank accession no.: OP933693.1) identified from fish in Lhasa River has similar genome organization with CsPV, and also has an unknown 145-aa-long protein at 3’ end, but no other published information available. So far, only the mentioned three fish parvoviruses belonging to unassigned genuses, subfamily *Parvovirinae* were characterized with uncommon genome structures. Moreover, CsPV is the first parvovirus identified from diseased marinculture fish, suggesting evolutionary adaptations to high salinity of CsPV and broadening the presence range of parvovirus. We speculated that it is necessary to elucidate the genetic diversity of fish parvoviruses and to explore the evolutionary relation. As with other fish papillomavirus, CsPaV contains the minimal backbone genes (E1, E2, L2 and L1), lacking any of the oncogenes (E5, E6 and E7) [9]. Of note, the L1 protein was expressed by a spliced mRNA, and a 67 bp segment with high AT content acted as an intron. Virus intron was rarely found in small viruses, while appeared in some viruses with large genome [37–39]. Besides, Lopez-Bueno et al. (2016) also reported that the SaPV1 L1 protein was expressed from a spliced transcript [23]. Although these are the only two evidences of the splicing events occurring in L1 encoding ORF in fish papillomavirus, we speculated that it may be a common strategy to infect fish by SaPV and CsPaV since they are both fish disease causing agents and co-infected with other viruses. And certainly, further studies are needed to address it.

Epidemiological investigations have shown that farmed Chinese tongue soles from Tianjin where the present study were conducted, Shandong and Hebei province in China were found to be positive for CsPV (ranging from 55%-100%) and CsPaV (100%). This indicated that an emerging disease may spread in the major Chinese tongue sole cultured area in China. Notably, wild Chinese tongue soles sampled from Bohai Sea showed positive for both viruses, and the L1 gene of CsPaV and VP gene of CsPV shared over 99% similarities with those from the wild ones, suggesting a possibility that CsPV and CsPaV may originated from wild Chinese tongue soles from Bohai Sea. Interestingly, researchers have identified parvovirus (isolate fi102par1, GenBank accession no.: OP933693.1) and papillomavirus (isolate fi100pap1, GenBank accession no.:OP933686.1) simultaneously in fish from Lhasa River by metagenomics in 2020, and no more information was published. Phylogenetic analyses showed that CsPV and CsPaV clustered with parvovirus of isolate fi102par1 and papillomavirus of isolate fi100pap1, respectively, implying a close evolutionary relationship between them. Researchers pointed out that wild fish may act as disease vectors of pathogens and transmit the pathogens to farmed fish [40,41]. However, direct evidence for viruses transmission through wild Chinese tongue soles in Bohai Sea, or even from fish in Lhasa River is lacking, and the origin of CsPaV and CsPV needs further study.

In conclusion, papillomavirus (CsPaV) and parvovirus (CsPV) were isolated and identified from the diseased Chinese tongue soles. Co-infection of the two viruses was responsible for the mass mortality of Chinese tongue soles. The two viruses appears to be emerging viral pathogens in farmed Chinese tongue soles. Further researches are needed to elucidate the mechanisms of virus co-infection, and methods should be conduct to prevent and control the emerging disease.

## Ethical statements

The study was performed in strict accordance with the Guide for the Care and Use of Laboratory Animals Monitoring Committee of Tianjin Normal University, China. The Chinese tongue soles were euthanized for 20–30 min in 1 mg/ml of MS-222 (Sigma, USA) before tissue collection.

## Supporting information

S1 Fig. Phylogenic tree was constructed basing on the nucleotide sequence of the L1 gene (http://pave.niaid.nih.gov).

S2 Fig. CaPaV and CsPV genome copies in intestine, muscle, heart, gill, mucus and eye tissues of experimentally infected fish at different time point (1, 2, 3, 5 and 7 dpi). The data represented means of three fish samples. The data were analyzed by Student’s *t* test (*P*<0.05, one asterisks; *P*<0.01, two asterisks).

## Acknowledgments

We greatly thank Dr. Xiao Zhang and Dr. Tong Hao for processing the sequencing data using bioinformatics platform.

## Author contributions

Conceptualization: Shuxia Xue, Jinsheng Sun

Data curation: Xinrui Liu, Yuru Liu, Shuxia Xue

Formal analysis: Xinrui Liu, Shuxia Xue

Funding acquisition: Jinsheng Sun, Lei Jia

Investigation: Siyu Yang, Zhu Zeng, Hui Li

Methodology: Xinrui Liu, Yuru Liu, Chang Lu

Resources: Lei Jia, Yanguang Yu, Houfu Liu

Validation: Xinrui Liu, Jiatong Qin, Yuxuan Wang

Writing-Original draft preparation: Shuxia Xue

Wring-Review and Editing: Shuxia Xue, Jinsheng Sun

